# Discrete Pain Behavior Without Systemic Inflammation in a Rat Osteotomy Model: A Platform for Analgesic Screening

**DOI:** 10.64898/2026.07.30.741923

**Authors:** Sahar Lotfollahi Soudmand, Shahabeddin Safi, Hamidreza Fattahian

## Abstract

**Background:** This study aimed to determine if a rat osteotomy model elicits a measurable systemic response and to utilize this profile to evaluate the mechanism of action of preemptive analgesics with translational relevance to veterinary perioperative pain management.

**Methods:** Twenty-five male rats were randomized into five groups: Sham, Surgery Control, Robenacoxib (2 mg/kg S.C.), Amantadine (30 mg/kg P.O.), and Combination. A femoral osteotomy was performed following ARRIVE 2.0 guidelines for refinement of surgical models. Serum levels of IL-6, PGE2, and cortisol were quantified via ELISA at baseline, 1-, 3-, and 6- hours post-surgery. Postoperative pain was assessed using the Rat Grimace Scale (RGS).

**Results:** The osteotomy model did not induce a significant systemic inflammatory or stress response. Serum IL-6 and cortisol levels showed no significant changes over time (IL-6: p=0.219; Cortisol: p=0.187) or between groups. While PGE2 showed a temporal increase (F=6.52, p=0.001), it was unaffected by drug treatments. In stark contrast, the model successfully produced significant pain-related behaviors in the Control group (p=0.007), which were effectively reduced by both Robenacoxib (p=0.014) and amantadine (p=0.019) monotherapies. The combination group also showed significant pain reduction compared to Control at T6 (p=0.002), with an additive effect relative to monotherapies. Correlation analysis confirmed a dissociation between systemic biomarker levels and pain scores.

**Conclusion:** The efficacy of Robenacoxib (a COX-2 inhibitor approved for veterinary use) and amantadine in the absence of altered systemic biomarkers suggests their analgesic actions are mediated predominantly through local or neurogenic pathways, with direct implications for optimizing perioperative analgesia protocols in companion animal orthopedic surgery.

## 1 Introduction

Deciphering the precise physiological mechanisms that drive postoperative pain behavior is a fundamental objective in behavioral neuroscience. Post-surgical pain arises from a complex integration of inflammatory, neurogenic, and neuropathic signaling pathways (1). To dissect these mechanisms, controlled animal models, such as rodent osteotomy, provide an essential experimental platform (2). While aspects of these models, including evoked behavioral responses (e.g., facial grimacing) and local neuro-immune mediators, have been characterized (3, 4), a critical question remains unanswered: Is the observed pain behavior in a controlled bone injury model coupled with a concurrent systemic physiological response? Specifically, the profile of the systemic inflammatory-stress axis—a key modulator of physiological state and behavior—in the acute phase following pure osteotomy has not been systematically defined, leaving a gap in the comprehensive physiological understanding of this model.

The body’s response to tissue injury caused by surgery can be conceptualized into three key and often overlapping frameworks: classic inflammation, which is the innate immune system’s response involving the release of mediators such as cytokines (like IL-6) and prostaglandins (like PGE2). These molecules contribute to the initiation and maintenance of hyperalgesia by directly activating and sensitizing peripheral nociceptors (5, 6). Alongside this response, neurogenic inflammation, which is triggered by the activation of nociceptors themselves and the release of neuropeptides such as Calcitonin Gene-Related Peptide (CGRP) and Substance P, plays a significant role in exacerbating local inflammation and transmitting pain signals (7). At the systemic level, a major stressor such as surgery activates the hypothalamic-pituitary-adrenal (HPA) axis, leading to the secretion of cortisol as the primary marker of the stress response. This hormonal response interacts in a complex manner with the systemic inflammatory response (8). The interaction of these local and systemic frameworks ultimately shapes the nociceptive behavior observed after injury.

Although previous investigations have delineated aspects of the local neuro-immune milieu in similar models (9, 10), the activation profile of the systemic inflammatory-stress axis in response to a pure osteotomy—especially during the critical acute phase—remains an uncharted component of the model’s integrated physiology. This gap limits our ability to interpret whether behavioral outputs, such as pain, are driven by local or systemic physiological states. The refinement of perioperative pain management in veterinary orthopedic surgery remains a critical welfare concern. Rodent osteotomy models serve as translational platforms for evaluating analgesic protocols prior to clinical application in companion animals (11). However, the validity of such models hinges on a precise characterization of their physiological profile—specifically, whether pain behavior correlates with systemic inflammatory markers that veterinarians routinely monitor (e.g., IL-6, cortisol). Without this foundational validation, extrapolation of preclinical findings to clinical veterinary practice risks misinterpretation of drug efficacy. The ARRIVE 2.0 guidelines emphasize that comprehensive physiological phenotyping is essential for model refinement and reproducibility in laboratory animal research (12). Robenacoxib, a highly selective COX-2 inhibitor, was selected due to its approved clinical use for perioperative pain management in companion animals (Onsior® in cats and dogs), making our findings directly translatable to veterinary practice. Amantadine, an NMDA receptor antagonist, was chosen because it targets central sensitization and wind-up phenomena, which are increasingly recognized as contributors to acute postoperative pain, particularly in orthopedic procedures where nociceptive input is intense and prolonged. Therefore, this study was designed to first map the systemic physiological response by measuring serum levels of IL-6, PGE2, and cortisol following osteotomy. Subsequently, and more pivotally, we aimed to leverage this systemic map as a diagnostic tool to interrogate the site and mechanism of analgesic action. We posited two mutually exclusive hypotheses: a significant systemic response would establish this model as a platform for testing broad anti-inflammatory drugs. Conversely, the absence of such a response—particularly if Robenacoxib shows efficacy in pain reduction without altering systemic markers—would strongly indicate that its primary analgesic mechanism is independent of systemic COX-2 inhibition, acting instead via local or neurogenic pathways. To test this, we employed Robenacoxib, a selective COX-2 inhibitor, and Amantadine, an NMDA receptor antagonist, not merely as treatments, but as pharmacological probes to dissect the physiological architecture of post-osteotomy pain behavior. This comprehensive physiological and behavioral assessment is essential for accurately defining the utility of this model and the mechanistic pathways of analgesic action within it.

## 2 Material and methods

### 2.1. Experimental Design and Subjects

A randomized, blinded experimental design was employed. Twenty-five adults male Wistar rats (250±50 g) were utilized in this investigation. The animals were sourced from the Laboratory Animal Breeding Center at the University of Tehran and acclimatized for two weeks prior to experimentation. Housing conditions included standard polypropylene cages maintained under controlled temperature (22±2°C), humidity (50±10%), and a 12-hour light/dark cycle with ad libitum access to standard pellet diet and water.

### 2.2. Ethical Approval

The experimental protocol received approval from the Research Ethics Committee of Islamic Azad University, Sciences and Research Branch (IR.IAU.SRB.REC.1404.010). All procedures were conducted in strict compliance with the National Institutes of Health (NIH) guidelines for laboratory animal care and use.

### 2.3. Study Groups

Animals were randomly distributed into five experimental groups (n=5 per group):

- Sham Group: Underwent anesthetic procedures and surgical preparation without osteotomy
- Control Group: Received osteotomy surgery without pharmacological intervention
- Robenacoxib Group: Administered 2 mg/kg subcutaneous Robenacoxib 30 minutes pre-operation
- Amantadine Group: Received 30 mg/kg oral Amantadine 30 minutes pre-operation
- Combination Group: Treated with both Robenacoxib (2 mg/kg S.C.) and Amantadine (30 mg/kg P.O.) 30 minutes pre-operation

### 2.4. Osteotomy Procedure

The surgical technique adhered strictly to the consensus standards for rodent bone healing models established by Histing et al. (11), with particular attention to minimizing soft tissue trauma to enhance model reproducibility and animal welfare—a core principle of the 3Rs. Following a 9- hour fasting period, general anesthesia was induced and maintained using Isoflurane (3-5% for induction, 1-2% for maintenance) delivered via precision vaporizer. The right femoral region was aseptically prepared, and a longitudinal incision provided surgical access. A unicortical defect (3 mm diameter) was created in the femoral diaphysis using a sterile low-speed orthopedic drill. Surgical closure involved 4-0 PGA absorbable sutures for muscular layers and 4-0 nylon for skin approximation. All surgical subjects received prophylactic cephazolin (20 mg/kg IM) immediately post-procedure, in accordance with established protocols for rodent orthopedic surgery (11). To preserve the accuracy of postoperative pain behavior assessment, no local anesthetics (e.g., bupivacaine splash or wound infiltration) were administered at any time during or after the surgical procedure

### 2.5. Behavioral Pain Assessment

Nociceptive behavior was evaluated using the Rat Grimace Scale (RGS) at predetermined intervals: baseline, 1, 3, and 6 hours post-operatively. A blinded observer analyzed 2-minute video recordings of facial expressions, scoring four distinct action units (orbital tightening, nose/cheek flattening, ear position, and whisker change) on a 0-2 scale. Cumulative RGS scores (maximum 8) were derived for each assessment point.

### 2-6. Blood Sampling and Analysis of Systemic Inflammatory and Stress Markers

To evaluate the systemic inflammatory and stress response, blood samples (approximately 300 µL per time point) were drawn from the lateral tail vein at four predetermined time points: baseline (T0, prior to any intervention), and at 1 (T1), 3 (T3), and 6 (T6) hours following the osteotomy procedure. Blood sampling was performed immediately after the 2-minute video recording for RGS assessment at each time point to prevent the stress of venipuncture from confounding the behavioral pain scores. Post-collection, blood was allowed to clot and then centrifuged at 1000 ×g for 15 minutes to separate the serum. The harvested serum was aliquoted and stored at -80°C to preserve analyte integrity until batch analysis.

#### 2-6-1. Interleukin-6 (IL-6)

The IL-6 concentrations were determined using the Rat IL-6 ELISA Kit (Karmania Pars Gene, Iran, Catalog No: KPG-RIL6). The assay is based on a sandwich ELISA principle. Briefly, standards and samples were added to the antibody-precoated microplate. After incubation and washing, a biotinylated detection antibody was added, followed by Streptavidin-HRP. Following another incubation and wash step, a chromogen substrate was added for color development, which was stopped before measuring the absorbance at 450 nm.

#### 2-6-2. Prostaglandin E2 (PGE2)

The PGE2 levels were measured using the Rat PGE2 ELISA Kit (ZellBio GmbH, Germany, Catalog No: ZB-10504C-R9648). This kit utilizes a biotin double antibody sandwich technology. Standards and samples were incubated in pre-coated wells, followed by the addition of a biotin-labeled detection antibody and Streptavidin-HRP. After incubation and washing, a chromogen solution was added. The reaction was stopped, and the absorbance was measured at 450 nm. The assay range is 50-1600 pg/mL with a sensitivity of 7 pg/mL.

#### 2-6-3. Cortisol level

Cortisol levels were assessed using a competitive ELISA. The specific protocol from the provided insert (IdeAL Diagnostic, Iran, Catalog No: 2924-96) was followed. In this method, cortisol in the sample competes with a cortisol-HRP conjugate for binding sites on biotinylated antibodies immobilized via streptavidin on the microplate. After incubation and washing, a chromogenic substrate (TMB) was added. The reaction was stopped, and the absorbance was read at 450 nm, with the intensity being inversely proportional to the cortisol concentration in the sample.

### 2-7. Euthanasia

Upon completion of the 6-hour post-operative monitoring and sample collection period, all animals were humanely euthanized. Euthanasia was performed by administering an intravenous overdose of thiopental sodium, ensuring a painless and ethical end point in accordance with animal welfare guidelines.

### 2.8. Statistical Analysis

Data were analyzed using SPSS 27. Normality was assessed via the Shapiro-Wilk test. A two-way repeated measures ANOVA analyzed the effects of Time, Group, and their interaction on systemic biomarkers (IL-6, PGE2, Cortisol). The non-parametric Friedman test assessed temporal changes in pain scores (RGS) within groups, and the Kruskal-Walli’s test compared groups at specific time points. Spearman’s correlation evaluated relationships between biomarkers and pain. Data are presented as Mean ± SD or median (IQR). A p-value < 0.05 was considered significant.

The sample size (n=5 per group) was chosen based on previous studies using the Rat Grimace Scale in surgical pain models, which demonstrated that this number is sufficient to detect significant differences in pain behavior (13, 14). This sample size is also consistent with the 3Rs principle of using the minimum number of animals required for reliable statistical inference, particularly given the large effect sizes observed in our primary outcomes

## 3 Results

### 3.1. The Systemic Inflammatory-Stress Response Profile Following Osteotomy

To determine whether the osteotomy model elicits a measurable systemic response, serum levels of key inflammatory mediators (IL-6 and PGE2) and a primary stress hormone (cortisol) were quantified at baseline and at 1-, 3-, and 6-hours post-surgery.

#### 3.1.1. Serum IL-6 Level

The experimental osteotomy procedure failed to induce a statistically significant systemic IL-6 response (Table1). A two-way repeated measures ANOVA revealed no significant main effect of time (F (2.20, 41.74) = 1.57, p = 0.219) and no significant interaction effect between time and treatment group (F (8.79, 41.74) = 0.57, p = 0.810) (Table2). Post-hoc analyses confirmed the absence of significant differences in IL-6 levels between any experimental groups at any time point (all P > 0.45). The mean IL-6 levels in the Surgery Control group were not significantly elevated compared to the Sham group at any post-operative interval.

#### 3.1.2. Serum PGE2 Levels

While a significant main effect of time on PGE2 levels was observed (F (3, 60) = 6.52, p = 0.001), with post-hoc tests confirming higher concentrations at T6 compared to all earlier time points (all P< 0.05), there was no significant main effect of group (F (4, 20) = 0.95, p = 0.46) and no significant time × group interaction (F (12, 60) = 1.45, p = 0.17) (**Table 2**). This indicates the temporal change in PGE2 was consistent across all groups and was not specifically driven by the osteotomy itself. Given that the Sham group also underwent the same anesthetic and surgical preparation (without osteotomy), the observed PGE2 elevation at T6 is most likely attributable to the physiological stress response to anesthesia, soft tissue manipulation, and the surgical procedure in general, rather than the bony defect specifically. Crucially, the selective COX-2 inhibitor Robenacoxib failed to significantly suppress this systemic PGE2 rise compared to the Surgery Control group, despite its clear efficacy in reducing pain behavior (**Table 1**).

**Table 1.** Serum levels of systemic inflammatory and stress biomarkers (Mean±SD) across experimental groups and time points.

| Group | Time | IL-6 (pg/mL) | PGE2 (pg/mL) | Cortisol (ng/mL) |
| --- | --- | --- | --- | --- |
| **Sham** | T0 | 27.74 ± 5.15 | 181.8 ± 16.74 | 108.0 ± 23.22 |
|  | T1 | 26.30 ± 4.76 | 179.9 ± 15.67 | 96.6 ± 25.98 |
|  | T3 | 29.68 ± 10.37 | 179.2 ± 23.37 | 98.4 ± 8.73 |
|  | T6 | 31.22 ± 6.58 | 227.2 ± 42.33 | 81.2 ± 23.47 |
| **Control** | T0 | 26.67 ± 5.83 | 195.2 ± 30.22 | 108.4 ± 24.03 |
|  | T1 | 24.72 ± 3.17 | 163.6 ± 36.61 | 100.0 ± 10.84 |
|  | T3 | 27.33 ± 4.75 | 194.8 ± 18.58 | 101.2 ± 6.87 |
|  | T6 | 28.76 ± 6.01 | 199.6 ± 42.99 | 98.4 ± 5.22 |
| **Robenacoxib** | T0 | 28.22 ± 2.45 | 189.2 ± 11.58 | 103.0 ± 3.08 |
|  | T1 | 28.78 ± 3.95 | 170.0 ± 25.80 | 104.6 ± 14.88 |
|  | T3 | 29.00 ± 6.44 | 196.0 ± 43.69 | 104.4 ± 1.67 |
|  | T6 | 27.29 ± 5.51 | 181.0 ± 53.15 | 97.8 ± 7.56 |
| **Amantadine** | T0 | 28.06 ± 0.61 | 188.2 ± 7.82 | 101.4 ± 2.97 |
|  | T1 | 28.60 ± 4.72 | 174.6 ± 18.31 | 98.0 ± 4.80 |
|  | T3 | 27.60 ± 8.93 | 199.2 ± 48.27 | 107.2 ± 6.14 |
|  | T6 | 28.10 ± 4.72 | 217.6 ± 40.70 | 91.2 ± 5.85 |
| **Combination** | T0 | 27.95 ± 0.77 | 179.0 ± 10.27 | 100.4 ± 1.52 |
|  | T1 | 28.20 ± 1.72 | 207.0 ± 33.02 | 106.4 ± 16.18 |
|  | T3 | 29.88 ± 1.33 | 187.4 ± 14.19 | 104.0 ± 7.18 |
|  | T6 | 33.55 ± 5.08 | 244.6 ± 26.29 | 116.2 ± 24.23 |

**Table 2.** Effects of time, treatment group, and their interaction on serum levels of systemic inflammatory and stress biomarkers (IL-6, PGE₂, and cortisol) as assessed by two-way repeated measures ANOVA.

| Biomarker | Effect | F-value | p-value | Partial $\eta^2$ | Statistical Power |
| --- | --- | --- | --- | --- | --- |
| **IL-6** | Time | F (2.20, 41.74) = 1.57 | 0.219 | 0.076 | 0.328 |
|  | Group | F (4, 20) = 0.57 | 0.810 | 0.107 | 0.237 |
| | Time × Group | F (8.79, 41.74) = 0.57 | 0.810 | 0.107 | 0.237 |
| **PGE2** | Time | F (3, 60) = 6.52 | 0.001 | 0.246 | 0.965 |
|  | Group | F (4, 20) = 0.95 | 0.460 | 0.160 | 0.257 |
| | Time × Group | F (12, 60) = 1.45 | 0.170 | 0.225 | 0.638 |
| **Cortisol** | Time | F (3, 60) = 1.65 | 0.187 | 0.076 | 0.412 |
|  | Group | F (4, 20) = 0.98 | 0.442 | 0.163 | 0.268 |
| | Time × Group | F (12, 60) = 1.56 | 0.128 | 0.238 | 0.651 |

Partial $\eta^2$ : (Partial Eta Squared) is an effect size measure indicating the proportion of total variance in the dependent variable (biomarker level) that is attributable to a given independent variable (e.g., Time or Group), after accounting for other effects in the model. Values of 0.01, 0.06, and 0.14 are conventionally interpreted as small, medium, and large effects, respectively (22). Statistical significance was defined as $P < 0.05$ . Degrees of freedom are reported as (numerator, denominator).

#### 3.1.3. Serum Cortisol Levels

No significant systemic stress response was detected. Analysis revealed no significant main effects of time (F (3, 60) = 1.65, p = 0.187) or group (F (4, 20) = 0.98, p = 0.442) on serum cortisol, and no significant interaction (F (12, 60) = 1.56, p = 0.128) (**Table 2**). Cortisol concentrations remained stable throughout the 6-hour post-operative period in all groups, as clearly demonstrated by the lack of a consistent increasing or decreasing trend across the experimental timeline in both the Sham and surgical groups (**Table 1**). Although the overall Kruskal-Wallis test at T6 showed a marginal significance (p=0.029), post-hoc pairwise comparisons revealed no significant differences between any specific groups (all adjusted p > 0.05), confirming the absence of a consistent treatment-related stress response.

### 3.2. Efficacy of Pharmacological Interventions on Pain Behavior

In stark contrast to the absent systemic biochemical response, the osteotomy model successfully induced significant pain-related behavior (Table 3). The RGS scores showed a significant increase over time in the Surgery Control group (p = 0.007). Both Robenacoxib (p = 0.014) and Amantadine (p = 0.019) monotherapies significantly altered the time-course of pain scores, demonstrating their analgesic efficacy. The combination therapy did not show a significant temporal effect (p = 0.106). Notably, at the 6-hour endpoint, all active treatment groups showed significantly lower pain scores compared to the Surgery Control (Table 5). Post-hoc pairwise comparisons with Bonferroni correction revealed that at T6, all active treatment groups had significantly lower RGS scores compared to the Surgery Control group (adjusted p < 0.05 for all comparisons); detailed pairwise comparisons at all time points are provided in Table 6. The combination group showed a progressive decline in RGS scores over time (T6: 0.23 ± 0.21), with significantly lower pain scores than the Control group at T6 (p=0.002, Kruskal-Wallis post-hoc), comparable to each monotherapy, suggesting an additive rather than synergistic effect

**Table 3.** Temporal evolution of postoperative pain behavior in experimental groups as assessed by the Rat Grimace Scale (RGS).

| Group | T0 | T1 | T3 | T6 | $\chi^2$ (Friedman) | p-value |
| --- | --- | --- | --- | --- | --- | --- |
| Sham | 0.03 ± 0.05 | 0.05 ± 0.09 | 0.08 ± 0.10 | 0.09 ± 0.06 | 2.76 | 0.431 |
| Control | 0.09 ± 0.02 | 1.81 ± 0.20 | 2.14 ± 0.14 | 2.10 ± 0.62 | 12.12 | 0.007 |
| Robenacoxib | 0.10 ± 0.02 | 0.88 ± 0.09 | 0.84 ± 0.17 | 0.76 ± 0.21 | 9.96 | 0.014 |
| Amantadine | 0.12 ± 0.01 | 0.76 ± 0.22 | 1.32 ± 0.32 | 0.70 ± 0.46 | 10.59 | 0.019 |
| Combination | 0.10 ± 0.02 | 0.42 ± 0.36 | 0.44 ± 0.33 | 0.23 ± 0.21 | 6.12 | 0.106 |

Data are presented as mean ± standard deviation (SD) of cumulative RGS scores (range: 0–8).
Pain behavior was evaluated at baseline (T<sub>0</sub>) and at 1 (T<sub>1</sub>), 3 (T<sub>3</sub>), and 6 (T<sub>6</sub>) hours post-surgery.
Within-group changes over time were analyzed using the Friedman non-parametric repeated measures test. Statistical significance was defined as $p < 0.05$ .

### 3.3. Disassociation Between Systemic Biomarkers and Pain Behavior

Spearman’s correlation analysis revealed a clear disconnection between systemic biomarkers and pain behavior (Table 4). No significant correlations were found between RGS scores and corresponding levels of IL-6, PGE2, or cortisol (all P> 0.05). The only strong correlations were between different time points of the RGS scores themselves (e.g., Grimace_T1 with Grimace_T3: rho = 0.892, P< 0.001). However, it should be noted that the small sample size (n=5 per group) may have limited the statistical power to detect weaker but potentially meaningful correlations.

**Table 4.** Spearman’s rank correlation coefficients (ρ) between serum biomarker levels and Rat Grimace Scale (RGS) pain scores at matched postoperative time points.

| Biomarker | Time Point | Correlation with RGS ($\rho$) | p-value | Interpretation |
| --- | --- | --- | --- | --- |
| IL-6 | T1 | 0.011 | 0.959 | No correlation |
|  | T3 | -0.041 | 0.848 | No correlation |
|  | T6 | 0.083 | 0.695 | No correlation |
| PGE2 | T1 | -0.114 | 0.588 | No correlation |
|  | T3 | 0.201 | 0.334 | No correlation |
|  | T6 | -0.440 | **0.028\*** | Weak to moderate negative correlation, of uncertain biological significance |
| Cortisol | T1 | -0.004 | 0.986 | No correlation |
|  | T3 | 0.253 | 0.222 | No correlation |
|  | T6 | -0.016 | 0.939 | No correlation |
| RGS (Internal) | T1 vs T3 | 0.892 | **<0.001\*\*** | Strong positive correlation |
|  | T1 vs T6 | 0.766 | **<0.001\*\*** | Strong positive correlation |

P < 0.05: weak to moderate negative correlation (P = -0.440) of uncertain biological significance.
\*\* P < 0.001: strong positive correlation between RGS scores at different time points, confirming the temporal consistency of pain behavior

### 3.4. Differential Effects of Pharmacological Interventions on Systemic Biomarkers and Pain

Kruskal-Wallis tests conducted at the 6-hour post-operative time point (T6) confirmed the absence of significant treatment effects on systemic inflammatory markers, revealing no differences between groups for IL-6 (P= 0.745) or PGE2 (P= 0.063) (Table 5). In stark contrast, the same analysis demonstrated robust and significant differences in pain scores between groups (P= 0.001). This clear dissociation underscores that the osteotomy model did not elicit a measurable systemic inflammatory or stress response; therefore, the pharmacological interventions had no systemic inflammatory state to modulate. Their efficacy in mitigating pain behavior, despite the absence of systemic biomarker changes, highlights the predominantly local or neurogenic nature of both the pain and the drugs’ mechanisms of action in this model.

**Table 5.** Between-group comparisons of systemic biomarkers and pain behavior at 6 hours post-surgery.

| Parameter |  | H-value | p-value<br>(Kruskal-Wallis) | Post-hoc Pairwise Comparisons at T6<br>(Bonferroni-adjusted p-values) |
| --- | --- | --- | --- | --- |
| **Systemic Biomarkers** | IL-6 | 1.951 | 0.745 | Not applicable (no significant main effect) |
|  | PGE2 | 8.925 | 0.063 | Not applicable (no significant main effect) |
| | Cortisol | 10.778 | 0.029 | Combination vs. Sham: $p = 0.073$ (marginally significant); other groups: $p > 0.05$ |
| **Pain Behavior** | RGS Score | 17.615 | 0.001 | Control vs. Robenacoxib: $p = 0.014$;<br>Control vs. Amantadine: $p = 0.019$;<br>Control vs. Combination: $p = 0.002$;<br>Treatment groups vs. each other: $p > 0.05$ |

**Table 6.** Pairwise comparisons of RGS scores between experimental groups at T1, T3, and T6 (Bonferroni-adjusted p-values)

| **Comparison** | **T1 (p-value)** | **T3 (p-value)** | **T6 (p-value)** |
| --- | --- | --- | --- |
| **Control vs. Sham** | 0.008 | 0.006 | 0.005 |
| **Control vs. Robenacoxib** | 0.042 | 0.038 | 0.014 |
| **Control vs. Amantadine** | 0.031 | 0.045 | 0.019 |
| **Control vs. Combination** | 0.021 | 0.019 | 0.002 |
| **Robenacoxib vs. Amantadine** | 0.412 | 0.389 | 0.451 |
| **Robenacoxib vs. Combination** | 0.335 | 0.298 | 0.287 |
| **Amantadine vs. Combination** | 0.421 | 0.402 | 0.356 |
| **Sham vs. Robenacoxib** | 0.289 | 0.312 | 0.298 |
| **Sham vs. Amantadine** | 0.305 | 0.298 | 0.287 |
| **Sham vs. Combination** | 0.278 | 0.265 | 0.243 |

## 4 Discussion

The dissociation between pain behavior and systemic biomarkers documented herein has direct implications for veterinary clinical practice. In companion animal orthopedic surgery, clinicians often rely on systemic markers (e.g., IL-6, cortisol) to guide analgesic protocols. Our findings caution against this practice in unicortical osteotomy models, suggesting that absence of systemic inflammation does not equate to absence of pain—a critical welfare consideration. Furthermore, robenacoxib’s efficacy in this model is particularly relevant as it is already approved for perioperative analgesia in cats and dogs (Onsior®). The demonstrated dissociation between its behavioral efficacy and lack of systemic PGE2 suppression suggests that robenacoxib’s primary site of action in orthopedic pain is local/peripheral rather than systemic—a finding that supports its safety profile in veterinary patients with comorbidities affecting systemic prostaglandin homeostasis.

The principal findings of this study demonstrate a clear dissociation between the neuro-behavioral output and systemic physiological state in a rat model of femoral osteotomy. Contrary to the initial hypothesis that a controlled bone injury would elicit a measurable systemic inflammatory-stress reaction, our results consistently showed that the surgical procedure, while effective in inducing significant pain-related behavior, failed to alter serum levels of IL-6 or cortisol, and produced a non-specific temporal increase in PGE2 that was refractory to COX-2 inhibition. Consequently, the primary analgesic actions of the preemptively administered drugs, Robenacoxib and Amantadine, appear to be independent of modulating the systemic inflammatory cascade, instead likely operating through local, peripheral, or neurogenic mechanisms.

The most striking finding of this study was the lack of a significant systemic IL-6 response following osteotomy. Numerous previous studies have consistently demonstrated that surgical procedures—ranging from abdominal surgery (15, 16) and thoracotomy (4) to intestinal manipulation (16) and incision models (11, 17)—induce a robust elevation of circulating IL-6 in rodents. This acute-phase cytokine response is widely considered a hallmark of surgical stress and tissue injury (8). In stark contrast, our femoral osteotomy model failed to elicit any measurable systemic IL-6 increase, even in the untreated surgical control group. This suggests that the femoral osteotomy, while sufficient to evoke robust pain behavior, may represent a localized insult that does not reach the threshold for initiating a profound systemic cytokine cascade. This could be attributable to the relatively limited soft tissue dissection and controlled nature of the bony defect, which might confine the inflammatory process to the local microenvironment—a concept well-supported by recent evidence indicating that immune responses in bone injury models are highly compartmentalized, with minimal systemic spillover (14). This notion is further supported by the concurrent lack of a systemic stress response, as indicated by stable serum cortisol levels. The stability of cortisol implies that the osteotomy procedure, under our experimental conditions, did not constitute a significant enough stressor to activate the HPA systemically (8). This collective absence of systemic IL-6 and cortisol elevation fundamentally redefines the pathophysiology of this specific osteotomy model, framing it predominantly as a model of local or neurogenic pain behavior, decoupled from systemic inflammatory-stress physiology.

In contrast to IL-6 and cortisol, a significant temporal increase in serum PGE2 was observed, peaking at the 6-hour post-operative mark. However, the absence of a significant Time×Group interaction, coupled with a similar upward trend in the Sham group, indicates that this rise was a non-specific phenomenon common to all animals subjected to prolonged anesthetic and peri-operative procedures, rather than a direct consequence of the osteotomy itself (8). The critical finding regarding PGE2 was the failure of the selective COX-2 inhibitor, Robenacoxib, to significantly suppress its systemic levels compared to the Surgery Control. This was unexpected, as the primary mechanism of action of NSAIDs is the inhibition of prostaglandin synthesis (18). This discrepancy can be attributed to several non-mutually exclusive mechanisms: (1) the systemic PGE2 pool measured in serum may originate primarily from constitutive COX-1 activity in platelets, vascular endothelium, and other tissues, which is insensitive to short-term COX-2 inhibition; (2) the administered dose of robenacoxib (2 mg/kg S.C.), while sufficient to achieve therapeutic concentrations at the local surgical site, may have been below the threshold required for significant systemic COX-2 suppression; and (3) the 6-hour observation window may have been insufficient to capture the pharmacokinetic accumulation needed for systemic effect. Conversely, had a non-selective COX inhibitor been employed, suppression of both COX-1 and COX-2 would likely have reduced systemic PGE2 levels, further supporting the notion that the circulating PGE2 pool in this model originates predominantly from COX-1 activity. This observation critically informs the interpretation of the drug’s analgesic action, strongly suggesting that its efficacy in reducing pain behavior is dissociated from a systemic anti-prostaglandin effect. This reinforces the model’s utility for distinguishing local from systemic drug pharmacodynamics The most compelling evidence from this study is the profound dissociation between the systemic physiological state and the evoked pain behavior. The osteotomy model robustly induced pain, as validated by significant increases in RGS scores in the Control group and their effective reduction by monotherapies, yet it failed to launch a concurrent systemic inflammatory-stress axis response. This decoupling is further corroborated by the lack of significant correlation between any systemic biomarker and pain scores. This paradigm forces a critical re-evaluation of the mechanisms of analgesic action in this model. The efficacy of both Robenacoxib and Amantadine in the absence of any measurable systemic anti-inflammatory or anti-stress effect strongly suggests that their primary site of action is not systemic. For Robenacoxib, its analgesic effect is likely mediated through potent local COX-2 inhibition at the site of injury, suppressing the release of pro-algesic prostaglandins in the surgical wound microenvironment without significantly impacting the systemic PGE2 pool. For Amantadine, its effect is presumably central, acting within the spinal cord or brain to attenuate central sensitization without modulating peripheral inflammation. The inefficacy of the combination therapy further suggests a potential pharmacodynamic interaction at these non-systemic sites, warranting further investigation. Collectively, these findings indicate that this model is exquisitely suited for screening analgesics that act via local or neural mechanisms, rather than broad-systemic anti-inflammatory drugs.

## Limitations of the Study

This study has some limitations. First, although our study lacked a positive pharmacological control (e.g., dexamethasone), prior work demonstrates that systemic anti-inflammatory agents can robustly suppress IL-6 in rat models of inflammation (19). Moreover, surgical interventions of moderate severity are known to reliably elevate plasma corticosterone in rats, a response that can even be attenuated by analgesics (20). The consistent absence of such biomarker changes in our osteotomy model—despite the use of validated ELISA assays capable of detecting physiological fluctuations in corticosterone (21)— strongly supports the conclusion that the behavioral output (pain) in this model is dissociated from a systemic inflammatory-stress physiology. Second, while our 6-hour observation window covers the typical peak for acute inflammatory and stress biomarkers, a delayed systemic response cannot be entirely ruled out. Third, our analysis focused on specific biomarkers (IL-6, PGE2, cortisol); measuring other key cytokines like TNF-α could have provided a more comprehensive picture. Finally, the sample size, while sufficient to detect large effects, may have been underpowered to identify more subtle changes.

## Conclusion

In conclusion, this study establishes the rat femoral osteotomy model as a refined platform for preclinical analgesic screening in veterinary orthopedics. The model evokes significant pain behavior—quantifiable via the validated Rat Grimace Scale—without triggering a confounding systemic inflammatory-stress response. This ‘clean’ physiological profile enhances the model’s specificity for evaluating local and neurogenic analgesic mechanisms. Critically, the efficacy of robenacoxib (a COX-2 inhibitor approved for veterinary use) and amantadine in this context supports their translational relevance for perioperative pain management in companion animals undergoing orthopedic procedures. Future studies should validate these findings in clinical veterinary settings while adhering to 3Rs principles for laboratory animal welfare.

## Acknowledgements

The authors sincerely thank Dr. Seyed Mohammad Sajjadi Dezfouli for his valuable contributions to the surgical procedures, anesthesia management, and pharmacological consultation. He has reviewed and consented to this acknowledgment

## Data Availability Statement

All raw data supporting the findings of this study, including individual RGS scores and serum biomarker measurements (IL-6, PGE2, and cortisol), are provided in S1 Data.

## Abbreviations

IL-6: Interlukin-6
RGS: Rat Grimace Scale
PGE2: Prostaglandin E2
CGRP: Calcitonin Gene-Related Peptide
HPA: hypothalamic-pituitary-adrenal axis
COX-2: Cyclooxygenase-2
NSAID: non-steroidal anti-inflammatory drugs
NMDA receptor: N-methyl-D- aspartate receptor

